# Pollen accumulation on hawkmoths varies substantially among moth-pollinated flowers

**DOI:** 10.1101/2022.07.15.500245

**Authors:** Gordon Smith, Christine Kim, Robert Raguso

## Abstract

Using the pollen loads carried by floral visitors to infer their floral visitation behavior is a powerful technique to explore the foraging of wild pollinators. Interpreting these pollen records, however, requires assumptions about the underlying pollen dynamics. To compare visitor foraging across flower species, the most important assumption is that pollen is picked up and retained on the visitor at similar rates. Given differences in pollen presentation traits such as grain number or stickiness even among flowers with similar morphologies, however, the generality of this assumption is unclear. We investigated pollen accumulation on the hawkmoth *Manduca sexta*, testing the degree to which accumulation differed among flower species and how pollen stickiness affected this accumulation. In no-choice floral visitation assays to six plant species visited by long-tongued hawkmoths in the wild, *M. sexta* individuals were allowed to visit flowers 1, 2, or 5 times, after which the pollen on their proboscises was removed and counted. We found that the six plant species varied orders of magnitude in the number of pollen grains deposited on the moths, with some placing thousands of grains after a single visit and other placing none after five. Plant species with sticky pollen adhesion mechanisms placed more pollen on the moths and had relatively less pollen accumulation over successive visits than non-sticky plants. Intriguingly, moths carried fewer pollen grains after 5 visits than after 2 visits, suggesting that both sticky and non-sticky pollen was lost during foraging. Together, our results suggest that interpretation of pollen load data should be made cautiously, especially when comparing across plant species.

## Introduction

Given the importance of ecosystem services provided by pollinators in both natural and agricultural settings (Ollerton, 2017), studying pollinator foraging behavior is critical to predicting their responses to both natural and anthropogenic perturbations. However, tracking the foraging decisions made by fast-flying floral visitors is challenging, especially in the natural floral communities in which these decisions are made. This is especially true for highly dispersive nocturnal pollinators such as hawkmoths (Sphingidae), due to rapid and unpredictable foraging paths that occur under dimly lit conditions (Baker, 1961; Martins & Johnson, 2007). While direct visit observations, interview choice assays (Campbell et al., 2016; Ogilvie & Thomson, 2015) and even camera traps at flowers (Edwards et al., 2015; Johnson et al., 2020) can provide key insights into their behavior, assessing the relative usage of different floral resources and specialization requires finer resolution tracking of complete foraging bouts.

One common method of inferring the foraging decisions of pollinators over longer time periods is to collect and identify the pollen carried on their bodies (Burkle et al., 2013; Scheper et al., 2014). As pollinators are unlikely to pick up a given plant’s pollen anywhere except from that plant’s flowers, pollen loads serve as a forensic record of which plants pollinators had visited during previous foraging bouts and can be collected from many more individuals than would be possible to observe. This technique is especially useful for studying the foraging of hawkmoths (Haber and Frankie 1989; Alarcón et al., 2008; Nilsson et al., 1987; Smith et al., 2021): as hawkmoths do not groom themselves or consume pollen, any pollen found on their proboscises is unlikely to be biased by decisions on which resources to collect (as seen in pollinators such as bees; e.g., Lunau et al., 2015). Drawing inferences beyond a given plant’s presence from these loads, however, requires several assumptions, the largest of which is that pollen from different flowers is acquired at relatively comparable rates.

While this assumption may be reasonable among closely related plants, per-visit pollen acquisition rates are likely to vary substantially across species based on a variety of plant traits. For example, plants vary orders of magnitude in the number and size of pollen grains they produce, from hundreds of 0.2-0.7mm long pollen grains in *Zostera* (Ruckelshaus, 1996) to many thousands of 10-12μm pollen grains in *Myosotis* spp. (Meudt, 2016). Furthermore, many plants display traits that improve pollen deposition and retention on their visitors, such as a sticky pollenkitt (Pacini & Hesse, 2005) or viscin threads (Hoch et al., 1993), that could significantly increase the number of pollen grains picked up during a single visit. Plants are also likely to differ in where they place pollen on their visitors based on floral dimensions such as nectar tube depth or stamen exsertion (Huang & Shi, 2013; Muchhala & Potts, 2007). For example, in flowers pollinated by Lepidoptera (butterflies and moths), the majority of pollen grains can be deposited anywhere from the proboscis (Bryant et al., 1991) to the face (Maad & Nilsson, 2004) to the wings (Cruden & Hermann-Parker, 1979; Murphy, 1984). Researchers, however, often collect pollen from only certain structures (e.g., the proboscis of hawkmoths or legs of bees), and may thereby miss grains deposited elsewhere even if some grains are detected at the collection site (Alarcon et al. 2008). Thus, while pollen loads may serve as a reliable record of *which* floral species had been visited, the degree to which pollen grain numbers can be used to draw inferences about the *relative frequency* of floral visitation remains unclear. This is especially true given that, to date, direct studies of pollen deposition on pollinators are rare (but see Harder & Thomson, 1989; Rademaker et al., 1997). Therefore, the degree of variation in deposition rates likely to be encountered in a floral community or pollinator guild is largely unknown, despite its importance in determining the inferences that can be drawn from this commonly used technique.

Guilds of hawkmoth-pollinated plants have evolved world-wide, showing convergent evolution for nocturnal anthesis, pale coloration, perfume-like aromas and deep nectar tubes or spurs (rev. by Johnson et al., 2017). However, because hawkmoth-pollinated plants often belong to very different angiosperm lineages, pollen morphology and pollen-placement mechanisms may differ substantively between species sharing hawkmoths as pollinators (Grant 1983; Haber & Frankie 1989). Here, we examine the number of pollen grains acquired by a representative long-tongued hawkmoth, Manduca sexta, on their proboscises over successive floral visits to six species of hawkmoth-pollinated plants with diverse floral morphologies and traits. We assess two key questions. 1) To what degree do plants differ in their pollen deposition on moth proboscises? Given the diversity of floral and pollen traits across the species studied, we predict that the plants should vary substantially in both the number of grains they deposit and in pollen accumulation over multiple visits by *M. sexta*. 2) How does a specific pollen trait – pollen stickiness – affect pollen deposition and pollen saturation on the proboscis? Here we present the answers to these questions, revealed through standardized, no-choice behavioral assays and blinded data analyses of the resulting pollen samples.

## Methods

### Study species

For this study we selected six night-blooming plant species that are visited by hawkmoths in nature and show diverse floral traits. *Datura wrightii* (Solanaceae) is a primary host plant and nectar resource for *Manduca sexta* in the southwestern USA (Bronstein et al., 2009), with large white flowers that moths completely enter to access nectar (Grant, 1983). *Ipomoea alba* (Convolvulaceae) produces similarly large and fragrant white flowers, and its copious nectar rewards are sought by hawkmoths in its native Argentina (Galetto and Bernardello 2004) and in naturalized, invasive populations elsewhere (Haber & Frankie, 1989; Johnson & Raguso, 2015). The nectar tubes (perianths) of *Mirabilis longiflora* (Nyctaginaceae) are similar in depth to those of *D. wrightii* and *I. alba*, and likewise are pollinated primarily by long-tongued hawkmoths (Grant and Grant 1983). In contrast, the related *M. jalapa*, a commonly cultivated plant, is typically visited and pollinated by smaller, shorter-tongued hawkmoths in its native range in Mexico (Martinez & Burquez, 1986). *Mandevilla macrosiphon* (Apocynaceae) shares this narrow tube morphology, and displays a highly specialized mechanism of pollen placement common to other *Mandevilla* species, whereby a sticky secretion is applied to their visitors’ long tongues before they contact the anther cones (de Araújo et al., 2014; Moré et al., 2007). Finally, *Oenothera harringtonii* (Onagraceae), endemic to Colorado, USA, is visited by pollen-robbing bees but is almost exclusively pollinated by hawkmoths (Skogen et al., 2016). Despite all target species being primarily pollinated by hawkmoths, these plants vary substantially in their floral dimensions (Figure 2) as well as other important floral traits. For example, both *O. harringtonii* and *M. macrosiphon* have mechanisms to adhere their pollen to visitors with sticky viscin threads (Hoch et al., 1993) and a sticky epidermal secretion respectively (de Araújo et al., 2014).

All plants used in this experiment were grown from seeds. Seeds for *Datura wrightii* and *Mirabilis longiflora* were collected from the Santa Rita Experimental range in Santa Cruz Co. in Arizona. Seeds of *Oenothera harringtonii* were collected from David Canyon, Comanche National Grassland, near La Junta, Otero Co. Colorado by K.A. Skogen, and seeds of *Mandevilla macrosiphon* were collected near Big Bend, Brewster Co. Texas by R.A. Levin. Seeds of *M. jalapa* and *I. alba* were acquired from commercial seed packets (Burpee, Inc.). Once sown, all plants were grown under greenhouse conditions in Sun-Gro Metro-Mix 360, with day/night temperatures of 24°C/21°C. Pressed vouchers were deposited for all plant species used at the L.H. Bailey Hortorium (BH), Cornell.

The floral dimensions of each species were measured from 10-15 individual flowers from at least 3 individual plants to the nearest mm using a metric ruler. For more methods details and the measurements for each plant species, see Supplementary Section [S1].

### Moth Rearing

Adult *Manduca sexta* moths used in this experiment were obtained from a laboratory colony maintained at Cornell. Larvae were reared in the lab on a cornmeal diet (Bell & Joachim, 1976; Goyret et al., 2009) on a long-day cycle (LD 16:8h; 24°C; 40-50% RH). Pupae were removed from the colony and isolated in a 31×31×32cm polypropylene mesh cage (BioQuip, Inc.) to eclose. Newly eclosed moths were kept under ambient conditions for 24-h prior to experimental flights.

### Flower visitation

At dusk, moths were moved from their holding cage into release tubes constructed from garden screening (Loew’s). While the moths acclimated, a single flowering plant was placed into a large (61×61×91 cm) polypropylene mesh cage (BioQuip, Inc.), oriented such that the flower (or flowers) were unobstructed. For *D. wrightii* plants, which were too large to fit into this cage, a single flower was fed into the cage while the rest of the plant remained outside. Once active, moths were released into the cage singly, and allowed to visit flowers until a) a given number of visits was reached, b) they had landed three times, or c) 10 minutes had elapsed. Moths were allowed to visit flowers 1, 2, or 5 times (N = 5 moths per visit number (3) for each plant species (6) = 90 total moth replicates). Within flights of a given plant species, the number of times each moth was allowed to visit flowers was chosen haphazardly, with a bias towards allowing more moth visits to ensure that the target sample sizes for 5-visit moths were reached. For representative videos of visits to each plant species, see Supplement [S2]. After each trial was completed, moths were re-captured and pollen on the proboscis (see Figure 1) was removed using ∼2 mm^2^ cubes of fuchsin gel (Kearns & Inouye, 1993), following the methods of Smith et al. (2021). These cubes were then melted onto clean microscope slides with a cover slip. All tools involved with producing slides were cleaned with alcohol swabs after each moth to prevent accidental pollen transfer.

**Figure 1:**
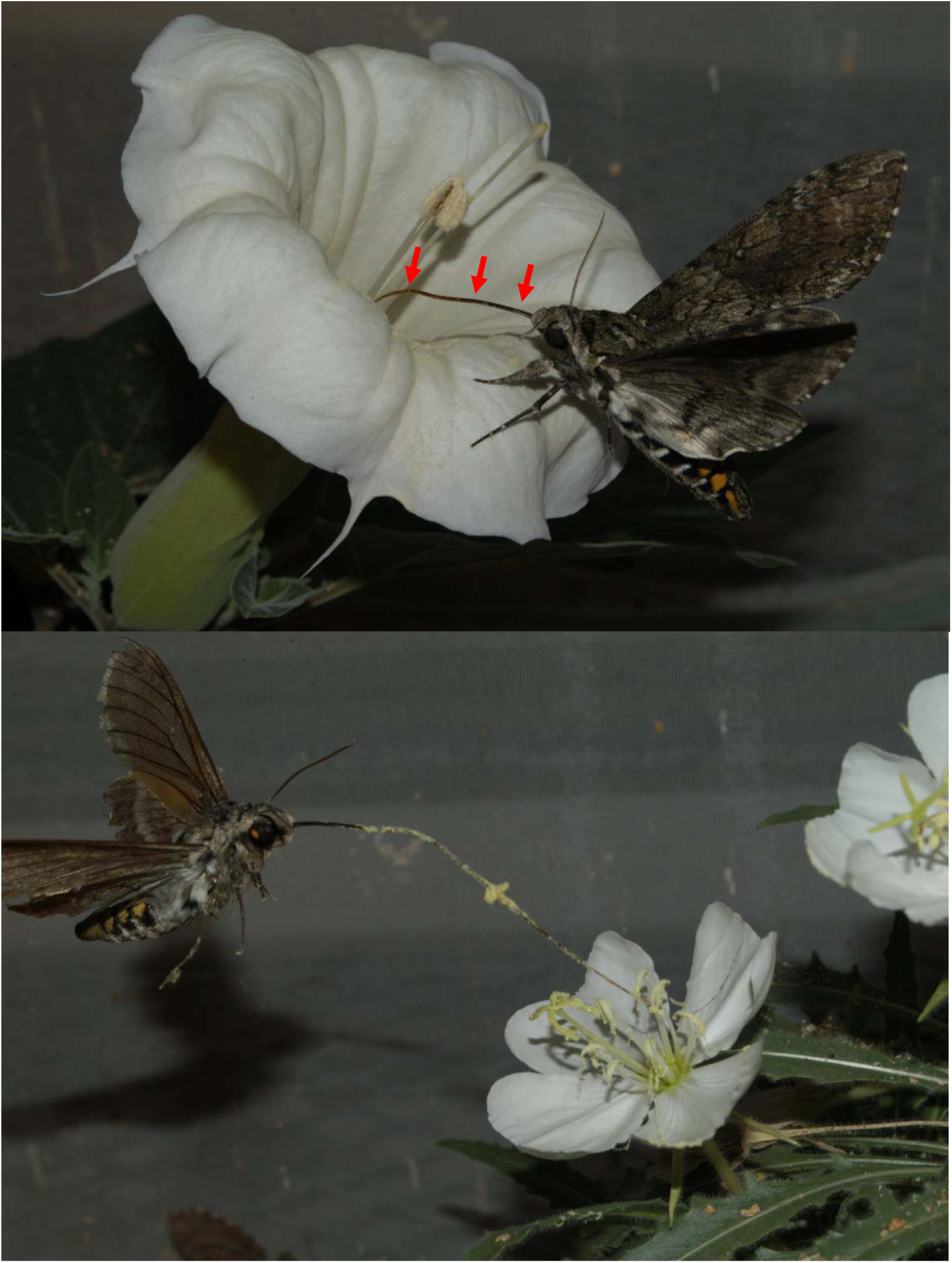
Pollen being deposited on the tongues of *Manduca sexta* while visiting *Datura wrightii* (top) and *Oenothera harringtonii* (bottom). While the yellow pollen of *O. harringtonii* is especially obvious, pollen can be seen on the proboscis of both moth individuals with the naked eye (highlighted with arrows on the moth visiting *D. wrightii*).

### Pollen counting

Pollen grains on slides from all species except *O. harringtonii* were individually counted at 10X-40X magnification using a light microscope (Nikon Eclipse 80i). For *Oenothera* pollen, which was too densely packed on the slides for individual grains to reliably be distinguished and counted, pollen numbers were estimated based on the size of the pollen clusters and their opacity, itself determined by the depth of the pollen in that cluster. For each cluster, the depth was determined by moving the microscope’s focal plane up and down to distinguish layers. Due to the flattened shape of the grains and variation in their orientation, clusters varied between 1.5 and 3.5 grains deep. Areas with similar opacity, and therefore depth, were measured with a scale and multiplied by the depth to estimate pollen grain numbers.

The total number of pollen grains presented by *Datura wrightii* and *Mandevilla macrosiphon* were estimated by vortexing fresh anthers in 70% ethanol, counting the grains contained in an aliquot of this mixture, and estimating the total number through multiplication. For *D. wrightii*, one anther (of 5 total) was vortexed in 200μL of ethanol, and grains were counted from 10μL. For *M. macrosiphon*, all anthers were vortexed in 100μL of ethanol, and grains were again counted from 10μL. The pollen presented by *Ipomoea alba, Oenothera harringtonii*, and *Mirabilis jalapa* were counted directly form fuchsin gel cubes that had removed all pollen from a subset of anthers, as the ethanol dilutions were not feasible for these species. For *O. harringtonii*, vortexing caused the viscin threads to clump pollen grains together rather than diluting them evenly. All *O. harringtonii* grains from one fresh anther (of 8 total) were removed with 2 gel cubes and were counted as described above. For *I. alba*, grains were removed from a pressed specimen; gel cubes were used to collect grains that had been dislodged from the anthers. For *M. jalapa*, the number of grains on fresh anthers was simply small enough that direct counting of every grain on a fuchsin gel cube was more accurate than estimates from dilutions. *Mirabilis longiflora* was not counted; all grains on pressed specimens were absent at the time of counting.

### Data analysis

All analyses were performed using R version 3.6.3 (R Core Team, 2020).

#### Question 1

To assess whether plant species differed in their pollen deposition on moth proboscises, we ran a Poisson generalized linear model (GLM) with pollen grain number as the response variable and plant species, visit treatment, and their interaction as fixed effects. Due to the small number of visit treatments and the fact that the numbers were not continuous (i.e., 3- and 4-visit treatments were not included), visit number was treated as a categorial variable rather than a continuous variable. For this model, moths that picked up 0 pollen grains were excluded; including these moths did not qualitatively change the results. To better examine the pollen acquisition pattern for each species, post-hoc models were run for each plant treatment independently. In these models, visit number was the only fixed effect, and all *p*-values were adjusted using a false discovery rate (FDR) correction.

#### Question 2

To assess whether pollen stickiness affected pollen deposition, we ran a Poisson GLM with pollen counts as the response and pollen stickiness, visit treatment and their interaction as fixed effects. For the purposes of this model, pollen stickiness was treated as a binary variable, with *M. macrosiphon* and *O. harringtonii* considered sticky and all other plant species considered not sticky.

## Results

### Pollen grains presented by flowers

The number of pollen grains presented to floral visitors differed substantially among plant species. In our greenhouse, each *Datura wrightii* flower produced an estimated 348,200 pollen grains, *Ipomoea alba* produced an estimated 1,625; *Mirabilis jalapa* produced an estimated 275; *Mandevilla macrosiphon* produced an estimated 1,100; and *Oenothera harringtonii* produced an estimated 10,968.

### Question 1: Variation in deposition

The number of pollen grains present on moth proboscises varied significantly, over three orders of magnitude, among plant species (GLM, see **Table 1, Figure 2**). Moths that visited flowers of *Oenothera harringtonii* carried the most pollen grains (mean grains ± SE: 1807.7 ± 297.4), followed by *D. wrightii* (mean 833.1 ± 185.9 grains), *M. macrosiphon* (452 ± 101.7 grains) and *I. alba* (12 ± 4.9 grains). Only one moth carried a single pollen grain of *Mirabilis jalapa* on its proboscis, and none of the moths that visited *M. longiflora* carried any pollen on their proboscis. The number of pollen grains also differed significantly between the visit treatments, with moths in the 2-visit and 5-visit treatments carrying more pollen grains than 1-visit moths across all plants. Visit number and plant species also interacted significantly, such that the shapes of the pollen accumulation curves for each species differed (**Figure 3**). In planned post-hoc analyses of all four plant species whose pollen was detected on moth proboscises, 2-visit moths carried more grains than either 1-visit or 5-visit moths (GLM *p* > 0.001 for *I. alba, D. wrightii, O. harringtonii* and *M. macrosiphon*). For D. *wrightii, O. harringtonii* and *M. macrosiphon* 1-visit moths carried fewer pollen grains that 5-visit moths; this pattern was reversed in *I. alba* (GLM p > 0.001 for all).

**Table 1:**
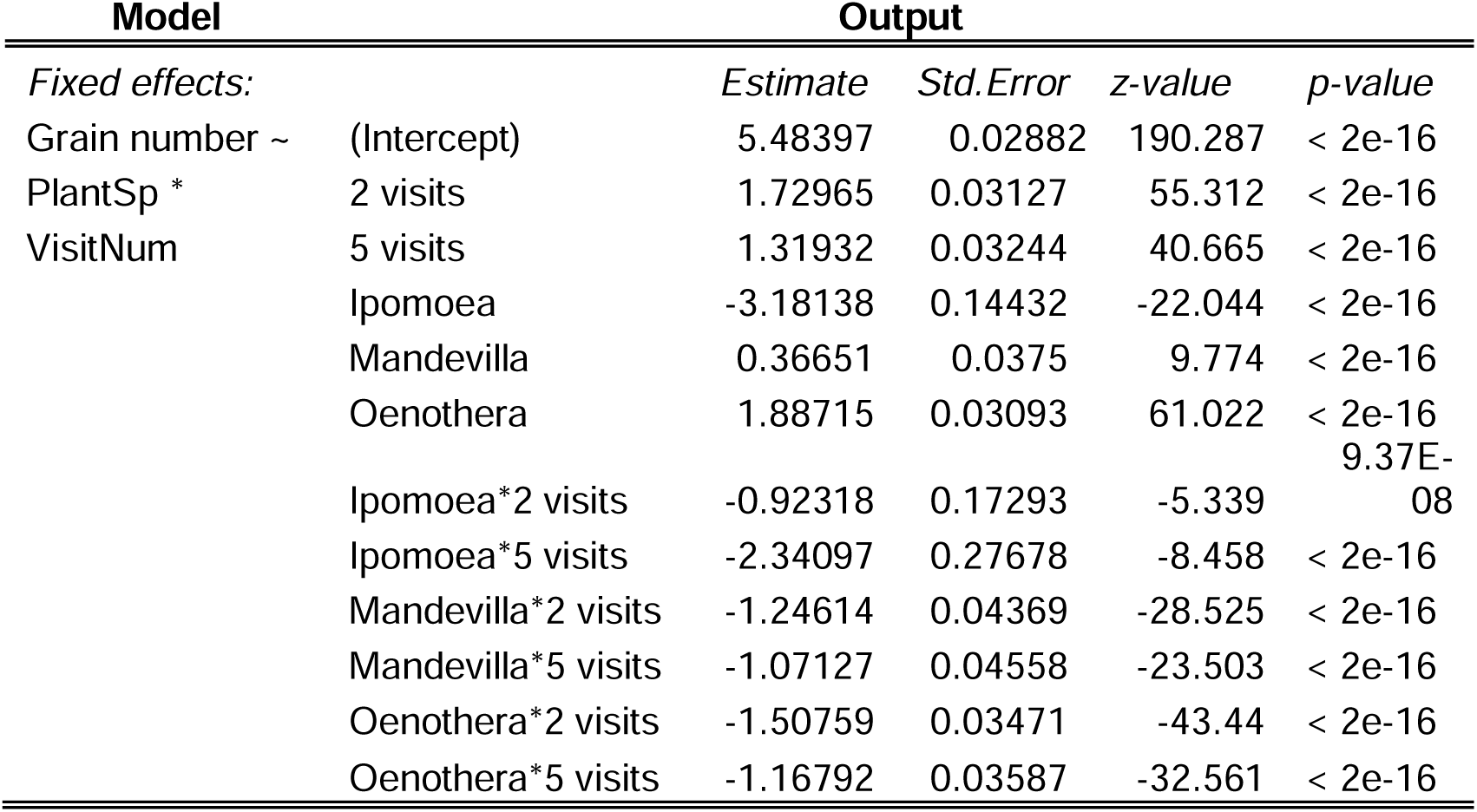
Model output for comparisons of pollen grain number across plant species.

**Figure 2:**
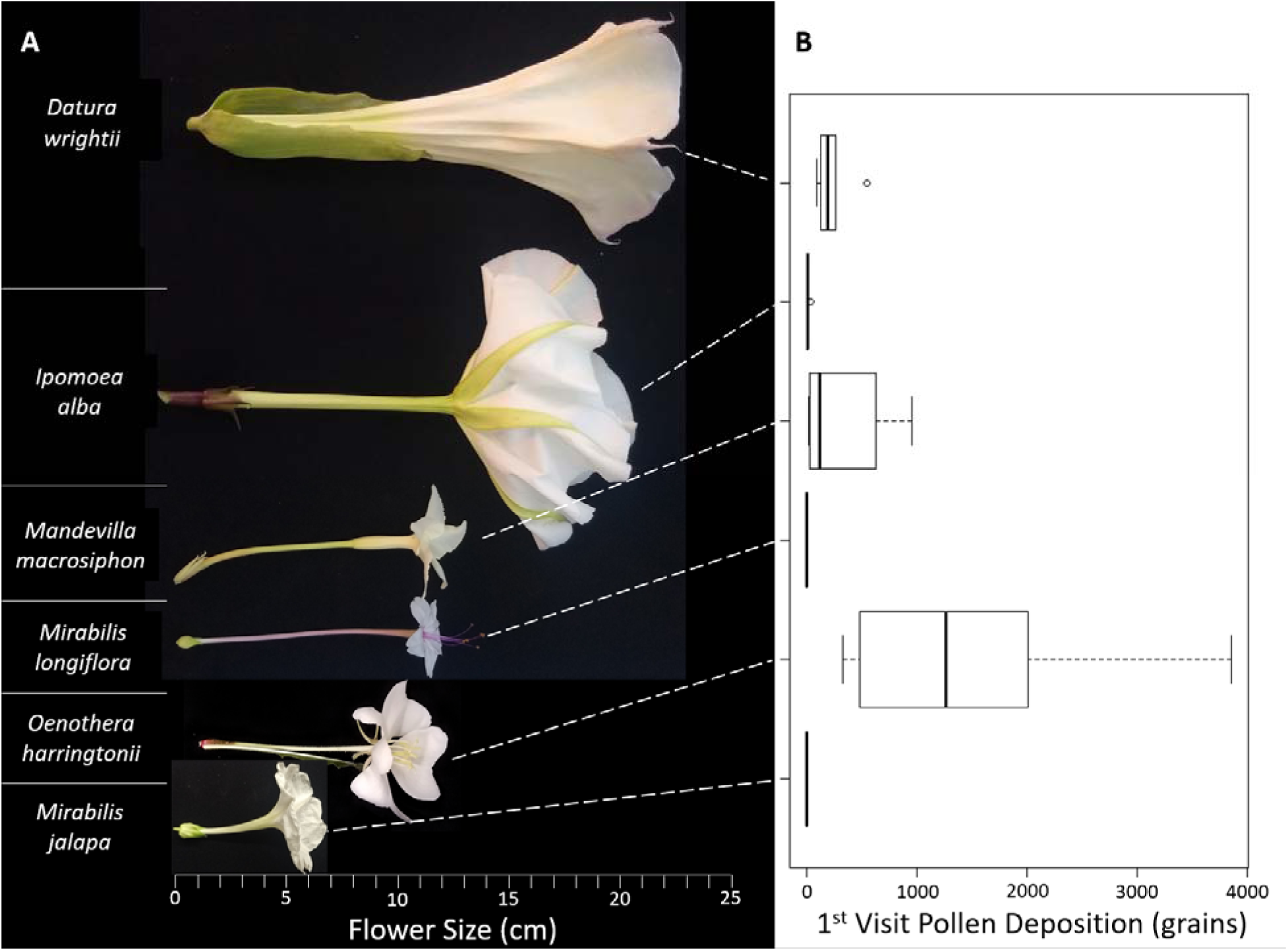
First-visit accumulation of pollen grains on the proboscises of *Manduca sexta* after visiting one of six plant species. *A*: Plant species arranged from largest to smallest based on nectar tube depth. *B*: Box-plot of pollen grain numbers present on moth proboscises after a single visit. The heavy bar represents the mean and the box bounds the 1^st^ and 3^rd^ quartiles.

**Figure 3:**
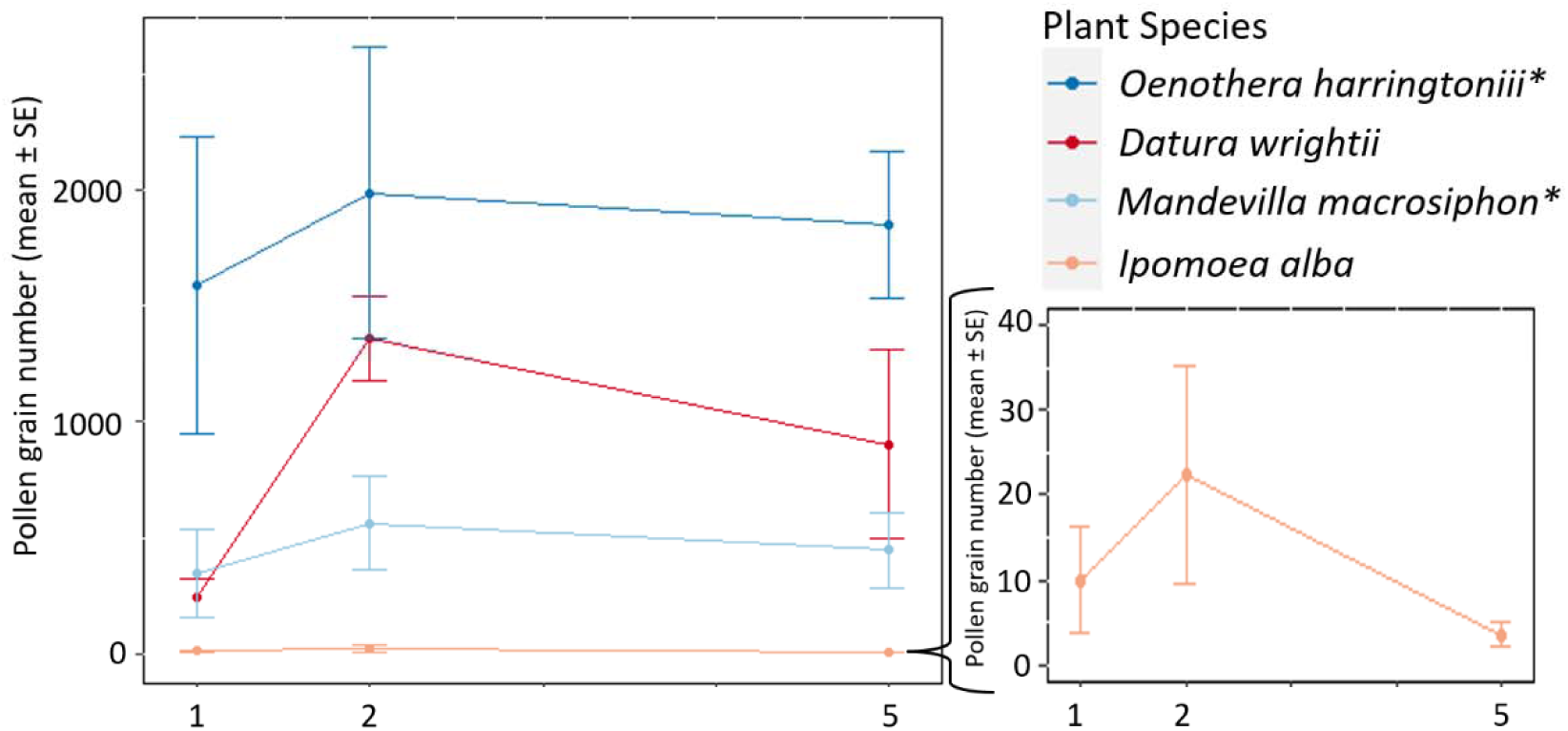
Accumulation of pollen on the proboscises of *Manduca sexta* moths by four plant species after 1, 2, or 5 successive floral visits. Due to the smaller number of pollen grains accumulated by moths visiting *I. alba*, data for this species is presented on both the main graph and the expansion below the legend. Plant species represented with blue and marked with an * had sticky pollen adhesion mechanisms.

### Question 2: the effect of pollen traits

Moths carried significantly more grains from plants with sticky pollen (**Table 2**). Pollen stickiness also interacted significantly with visit treatment such that 1-visit moths carried relatively more grains compared with 2-visit and 5-visit moths in sticky plants. Thus, the relative differences between the visit treatments were smaller in sticky plants (**Figure 3**).

**Table 2:**
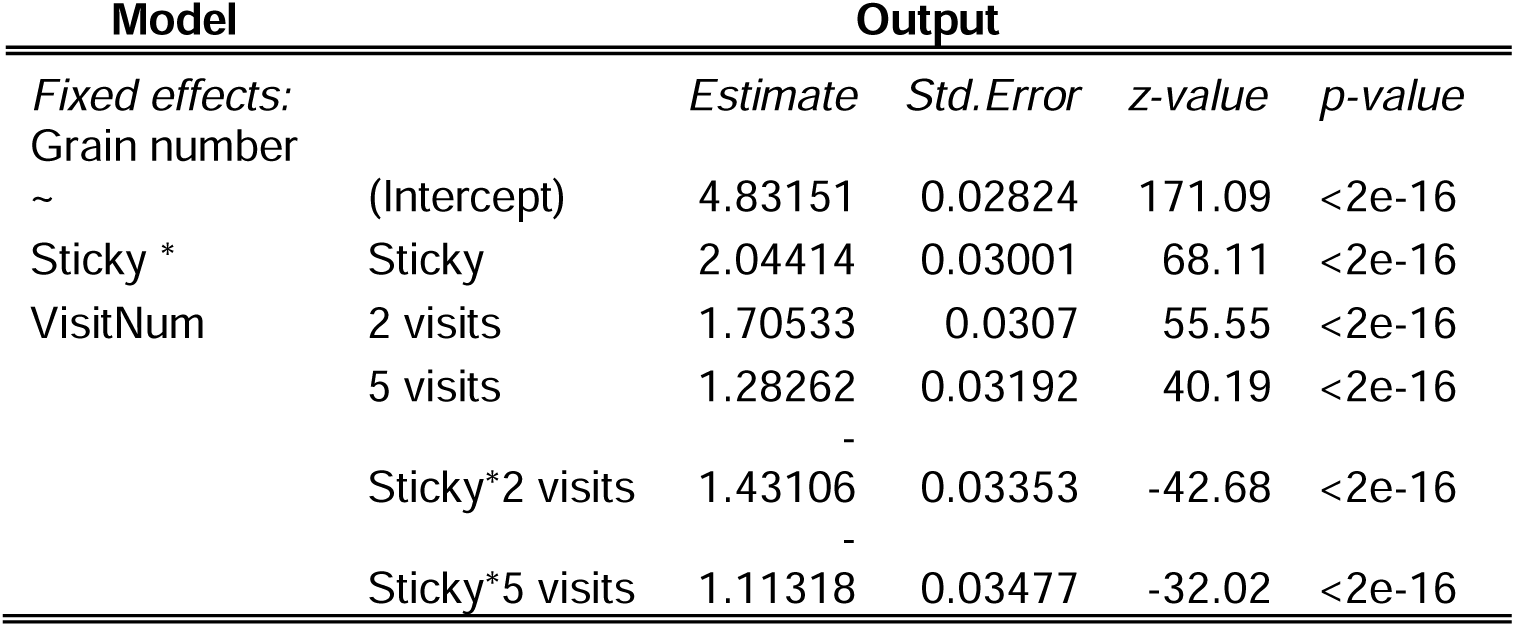
Model output for comparisons of pollen grain number between plants with sticky and non-sticky pollen

## Discussion

Studying the pollen loads of foraging pollinators can allow the examination of many complete foraging records, but interpreting those records relies on several important assumptions. Here, we tested one such assumption by examining the degree to which the number of pollen grains picked up by the moths are likely to vary between plant species. We found that the plant species tested varied orders of magnitude in the number of pollen grains they placed on the moths’ proboscises, that the shape of accumulation curves likewise varied between plants, and that plant traits such as pollen stickiness can have large impacts on the pickup and retention of pollen grains.

Together, these results clearly suggest that the number of pollen grains detected in a pollen load is a poor predictor of the number of visits or foraging effort allocated by an individual forager towards a given flower species. In some cases, even the presence or absence of pollen may be deceiving, as neither of the *Mirabilis* species we tested deposited almost any pollen grains on moth proboscises even after five visits. This absence of grains was likely due at least in part to the pollen of these species primarily contacting other sites on their visitors, such as the highly extruded anthers of *M. longiflora* (Supplement S1) contacting the head and eyes of our moths. However, moth proboscises did contact the anthers of these species during probing and nectaring, leaving the possibility that the large, smooth pollen grains simply adhered poorly to that structure despite prior reports of pollen from this species being found on the proboscis (Alarcón et al., 2008; Grant & Grant, 1983). Collecting pollen from other areas on the moth’s body may help alleviate this issue, when feasible (Moré et al., 2006).

It is also clear, and unsurprising, that pollen traits affect both the number of pollen grains deposited and the accumulation of grains over subsequent visits. In particular, the viscin threads of *Oenothera* and sticky floral secretion of *Mandevilla* were correlated with higher pollen grain numbers than the less sticky *Ipomoea* and *Datura*. This effect was most apparent in the smaller differences between the 1-visit and 2-visit treatments for *O. harringtonii* and *M. macrosiphon*, which suggests that a single visit was nearly sufficient to saturate the moth’s proboscis with pollen. Such high efficiency pollen transfer would be especially valuable for plant species that do not occur in highly dense populations, either due to their growth form and life history or to habitat fragmentation (see Suzan et al., 1994). These species also placed a large proportion of the total grains they produced on their visitors: a single visit removed ∼15% and ∼40% of total pollen from *Oenothera* and *Mandevilla*, respectively. While not quite as efficient as plants with pollinia or other pollen packaging mechanisms (e.g., orchids, milkweeds; Nilsson, 1983; Woodson, 1954) where a single pollinator visit can remove a pollinium containing thousands of pollen grains, these high removal proportions suggest that sticky pollen may reduce both pollen discounting rates and the potential missed mating opportunities associated with producing less pollen.

Pollen grain production is likely a large determinant as well: in addition to being sticky, our *Oenothera harringtonii* flowers produced ∼11000 pollen grains and therefore had many more grains available for transfer than *Mirabilis jalapa*, which produced less than 300. While the total amount of *M. longiflora* pollen available was not directly counted in this study, it likely produced a similar numbers to its congener (∼150-800, based on pollen:ovule ratios and a single ovule in this genus; Cruden, 1973). These low numbers likely interacted with other factors such as placement location and pollen morphology, resulting in the low (or absent) pollen grains moths picked up from these species. The high number of pollen grains produced by *Datura wrightii* (∼350,000) was also likely a major contributor to the fact that *D. wrightii* species placed the 2^nd^ highest number of grains on moth proboscises without being especially sticky.

In addition to assumptions about pollen acquisition, interpreting pollen loads often assumes that once pollen is acquired, it is not lost. Intriguingly, our data also cast doubt on this assumption, as all the plant species that placed any pollen on the moths’ proboscises saw declines in the number of grains present between the 2^nd^ and 5^th^ visit. For example, 5-visit moths visiting *Datura* carried ∼66% of the pollen carried by 2-visit moths, and on *I. alba* 5-visit moths carried only ∼16% of the grains carried by 2-visit moths. These patterns most likely suggest that pollen is being lost from the proboscis between the 2^nd^ and 5^th^ visit. While the specific causes of this loss are not clear from this study, we speculate that the majority is due to passive loss (Inouye et al., 1994) resulting from proboscis curling. In hawkmoths, curling the proboscis after feeding can result in substantial pollen movement and loss from this structure (Smith et al. 2021). Anecdotally, repeated curling events were more likely to have occurred for 5-visit moths: after 1-3 visits to focal flowers in quick succession (during which there may not have been even 1 proboscis curl), moths often explored the rest of the cage and examined other parts of the plant before returning for the 4 and 5 visit. Intriguingly, while the loss was somewhat dampened in *O. harringtonii* and *M. macrosiphon*, the fact that that even sticky pollen was lost suggests that loss due to proboscis curling may be a relatively general phenomenon for plants placing loose pollen (rather than pollinia) on hawkmoth proboscises.

### Conclusions

Our results suggest that pollen accumulation on hawkmoth proboscises is highly variable, even across diverse flowers presumably adapted to hawkmoth pollination, and therefore that comparisons across plant species should be made with care. This is not to say, however, that grain numbers in pollen loads do not provide valuable information. For example, comparisons *within* species may be more reliable: for most of our plants, the degree of variation within across treatments (visit number) within species was relatively low compared with the differences among plant species. Thus, while determining the relative foraging effort of single wild-caught moths on *Datura* versus *Oenothera* may not be possible based on pollen grain numbers, it may be possible to compare the number of *Datura* grains carried by two different individuals to explore a number of questions, such as their relative foraging effort to that plant or the impacts of variation in tongue length on pollen transfer. Further examination of these patterns of pollen accumulation and loss from floral visitors would be valuable in more accurately interpreting animal-carried pollen loads.

## Notes

### Competing Interest Statement

The authors have declared no competing interest.

